# Maximally Divergent Synonymous Gene Design with SIRIUS

**DOI:** 10.64898/2026.04.06.716428

**Authors:** Amirsadra Mohseni, Ian Wheeldon, Stefano Lonardi

## Abstract

The design of maximally divergent DNA sequences translating into the same protein is a critical problem in synthetic biology. Current design tools that rely on heuristics or machine learning often fail to effectively minimize the length of shared subsequences between the gene copies, compromising strain stability. Here, we introduce SIRIUS, a combinatorial optimization algorithm designed to generate maximally divergent coding sequences for a given protein of interest. Leveraging integer linear programming enforcing host-specific codon usage thresholds, SIRIUS stabilizes synthetic constructs and broadens the accessible design space for robust and scalable synethtic biology. Experimental results show that SIRIUS produces diverse sequences with fewer shared subsequences than existing methods. SIRIUS is freely available on GitHub at https://github.com/ucrbioinfo/sirius.

## Introduction

In synthetic biology, it is often necessary to integrate multiple copies of the same gene into the genome of unicellular organisms to increase the production of a particular protein of interest. However, introducing identical DNA copies can have drastic consequences: when these copies share sufficiently long identical subsequences, recombination events may occur, leading to gene loss and reduced strain stability (see, e.g., [1, 2, 3, 4, 5]). To minimize this problem, one needs to introduce as many sequence variations in each copy of the gene as possible. In order to maximize the difference (or divergence) between the copies, one can rely on the redundancy of the genetic code to diversify the codon usage for the same amino acid for the protein of interest. Since most amino acids are encoded by two to six synonymous codons, the number of genes that translate into the same protein is exponential relative to the length of the protein. Given the combinatorial explosion of the search space, the problem of designing maximally divergent genes is computationally challenging. The ability to generate sets of synonymous gene sequences that are both codon-optimized and mutually divergent is essential for advancing strain engineering, ensuring construct stability, and supporting industrial-scale protein production. This motivates the need for design tools that can efficiently explore the vast search space and synthesize maximally divergent, codon-constrained sequences.

In this work, we formulate the *synonymous gene design* problem as an integer linear program (ILP). In this combinatorial optimization framework, the input is (a) the sequence of amino acids for the protein of interest *P*, (b) the desired number of copies *N* for the genes encoding *P*, and (c) an optional species-specific codon usage table. The objective is to generate *N* synonymous genes that translate into *P* and minimize the number and frequency of common subsequences over all pairs of synonymous genes. The problem of solving ILPs optimally has been extensively studied in computer science and is known to be NP-hard [6].

Several variants of the synonymous gene design problem have been studied, and software tools have been developed for this purpose. For instance, Fallahpour *et al*. introduced CodonTransformer, a transformer-based model trained on multiple organisms to generate divergent degenerate DNA sequences [7]. CodonTransformer, however, does not accept user-defined codon usage tables without retraining, and it requires the use of codon bias. CodonBERT by Ren *et al*. is another transformer-based approach that produces a single codon-optimized sequence from a given amino acid input [8]. Hossain *et al*. developed the Non-Repetitive Parts Calculator (NRPC), a broader framework that includes synonymous gene design among many other functions. Similarily, Tuller *et al*. developed ChimeraUGEM, a multi-objective evolutionary algorithm, to reduce the number of homologous subsequences between genes [9]. Both NRPC and ChimeraUGEM, however, do not minimize shared subsequences directly; instead, they require a user-specified maximum subsequence length *l*_*max*_, which limits the length of the longest common subsequence for all pairs of gene copies [10]. In these tools, if a set of *N* gene copies that can satisfy the threshold *l*_*max*_ does not exist, the algorithms returns a smaller number of copies that adhere to the specified *l*_*max*_. Diez *et al*. created iCodon, which employs a genetic algorithm to optimize codon usage for a given amino acid sequence [11]. iCodon, however, neither minimizes long shared subsequences nor extends beyond a limited set of model organisms. Among existing methods, the only tool that directly addresses the synonymous gene design problem is GeneDiversifier [12], which employs a greedy strategy to reduce long shared subsequences through local clustering. However, GeneDiversifier does not ensure that the solution will be optimal.

In response to these needs, we present SIRIUS (*Systematic Identification of Redundant Identically Translated Sequences*^*^), an efficient algorithm that can generate maximally divergent codon-optimized DNA sequences for a given protein of interest. SIRIUS leverages combinatorial optimization to systematically explore the vast search space of synonymous codon choices while enforcing host-specific codon usage thresholds and minimizing long shared subsequences across output sequences. We demonstrate that SIRIUS consistently produces sets of synonymous genes with fewer shared subsequences than existing tools, including GeneDiversifier, on a panel of therapeutically and industrially relevant genes. By improving sequence divergence without compromising codon optimization, SIRIUS enables the stable multi-copy expression of genes in biomanufacturing strains, addressing a longstanding bottleneck in strain engineering.

## Methods

At a high level, SIRIUS proceeds as follows. First, the input protein *P* is expanded into all possible codon choices for each amino acid. Next, SIRIUS constructs an ILP where (1) base variables represent codon and nucleotide assignments, (2) additional variables model the presence of common subsequences, and (3) the objective function minimizes both the length and the number of such common subsequences across all sequence pairs. Finally, SIRIUS leverages the Google OR-Tools CP-SAT solver to solve the ILP [13]. The optimal solution of the ILP represents the most divergent set of *N* genes that translates into protein *P* and optionally satisfy user-defined codon usage constraints. An overview of this workflow is shown in Figure 1. In Step 3, common subsequences between sequence pairs are highlighted. Some common subsequences, however, are unavoidable. For example, all codons encoding alanine (`A`) begin with `GC`; therefore, when alanine follows methionine (`M`), every pair of genes must share a homologous subsequence of length 5.

**Figure 1:**
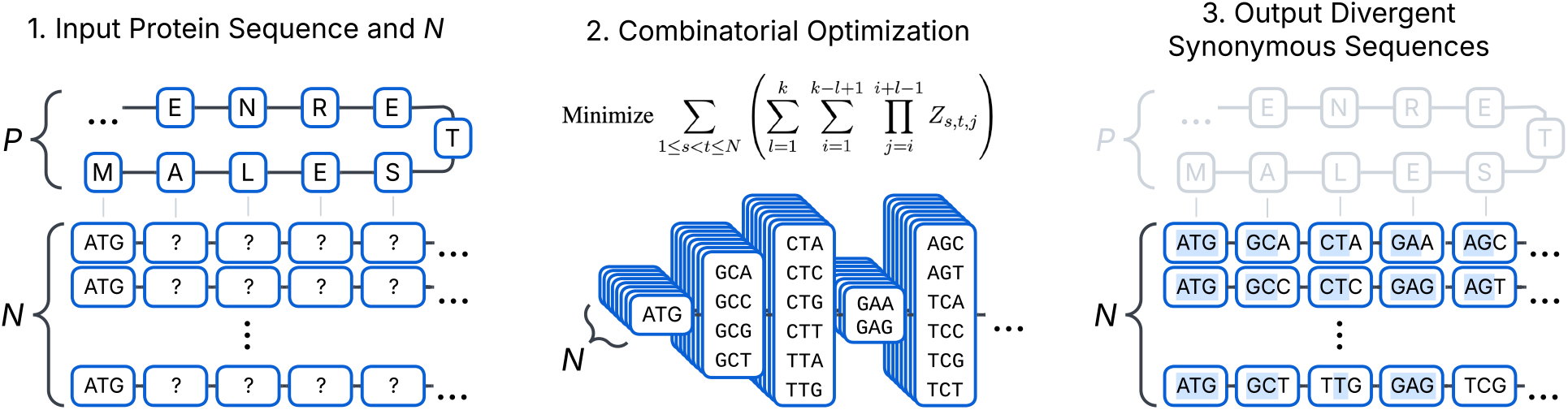
Overview of the SIRIUS workflow. **Step (1)** The input to SIRIUS is a protein sequence of interest *P* and the desired number *N* of synonymous DNA sequences to be designed; **Step (2)** SIRIUS solves an integer linear program with the objective function shown here, with potentially millions of variables and constraints; below the objective function the figure illustrates the codon choices for each amino acid in the example peptide *P* ; **Step (3)** SIRIUS produces of *N* synonymous DNA sequences that encode *P* with the fewest and shortest common subsequences; the light blue highlights indicate common subsequences between any pair.

### Problem Definition

The problem we address here is as follows. Given a target protein *P* and the number *N* of desired synonymous genes, we want to design a set *S* of *N* gene sequences such that (i) all the genes in *S* translate into *P*, and (ii) the length and the number of common subsequences between any pair of genes in *S* are minimized. We formalize this objective as follows. For each pair of aligned genes *i* and *j* in *S*, we identify the set of maximal common subsequences between *i* and *j*. Then, we count the number of common subsequences of each length over all pairs of genes. We define *D*(*S*) = U_1≤*l*≤3|*P*|_{(*l, n*_*l*_(*S*))} as the set of (length,count) pairs for *S*, where *n*_*l*_(*S*) is the number of common subsequences of length *l* (if *n*_*l*_(*S*) = 0, the pair is not included in *D*(*S*)). To determine whether solution *S*_1_ is better than solution *S*_2_, we compare *D*(*S*_1_) and *D*(*S*_2_) lexicographically, i.e., we start from the longest common subsequence length *l* in either *S*_1_ or *S*_2_. If *S*_1_ has fewer common subsequences of length *l* than *S*_2_, then *S*_1_ is better. If *S*_2_ has fewer common subsequences of length *l* than *S*_1_, then *S*_2_ is better. If *S*_1_ and *S*_2_ have the same number of common subsequences of length *l*, we proceed to the next lower length and repeat the same process. The pseudo-code for this procedure is provided in the **Supplementary Notes**.

#### Example 1.

Let *S*_1_ and *S*_2_ be two solutions. Let us assume that *D*(*S*_1_) = {(3, 1), (2, 5), (1, 5)} and *D*(*S*_2_) = {(3, 0), (2, 6), (1, 7)}. In solution *S*_1_, there is one shared subsequence of length 3, five subsequences of length 2, and five subsequences of length 1. *S*_2_ is a better solution than *S*_1_ because *S*_2_ has fewer shared subsequences of length 3, even though *S*_2_ has more shared subsequences of length 2 and length 1.

#### Example 2.

Let *S*_1_ and *S*_2_ be two solutions. Let us assume that *D*(*S*_1_) = {(5, 1), (3, 7), (2, 5)} and *D*(*S*_2_) = {(5, 1), (3, 5), (2, 7)}. In this case, both solution *S*_1_ and solution *S*_2_ have the same number of common subsequences of length 5. *S*_2_ is a better solution than *S*_1_ because *S*_2_ has fewer subsequences of length 3, even though it contains more shared subsequences of length 2.

We formalize the problem as follows. Let *n* ∈ {1, 2, …, *N* } be the input sequence index, and *p* ∈ {1, 2, …, |*P*|} be the amino acid position in protein *P* (where *P* is the length of *P*). We use *P* [*p*] to denote the *p*-th amino acid in *P* (e.g., if *P* starts with a methionine, then *P* [1] = `M`). We encode the mapping between codons and amino acids using table *T* . Given an amino acid *a*, we define *T* [*a*] to be the list of codons for *a* (e.g., for valine, we have *T* [`V`] = {`GTT`, `GTA`, `GTC`, `GTG`}). We use *c* ∈ {1, 2, …, |*T* [*a*]|} as the codon index for amino acid *a*, and *b* ∈ {1, 2, 3} as the base position within codon *c*.

### Integer Linear Programming Formulation

We formulate an ILP in which there is one binary variable *x* for every base *b* of every codon option *c* of every amino acid position *p* of every gene copy *n*. We set *x*_*n,p,c,b*_ = 1 when base *b* in codon *c* of amino acid position *p* in sequence *n* is chosen, and 0 otherwise. Henceforth, for clarity, we will overload the notation for *c* to also represent a codon itself.

For each codon option, we introduce an auxiliary binary variable *y* that is equal to 1 when all of its composing bases are 1. Formally, we set *y*_*n,p,c*_ = 1 if codon *c* is selected for amino acid position *p* in sequence *n*. In order for a codon *c* to be chosen, we require that all of its composing base variables also be chosen. We establish the dependency between base variables and codons using the following constraint

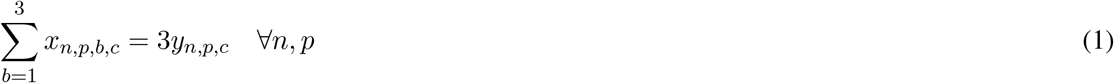

which implies that *y*_*n,p,c*_ = 1 if all bases of codon *c* are 1 (i.e., all *x*_*n,p,b,c*_ = 1), and *y*_*n,p,c*_ = 0 if all bases of codon *c* are 0 (i.e., all *x*_*n,p,b,c*_ = 0). No other combination for *x*_*n,p,b,c*_ is allowed (i.e., this equation does not allow some bases of a codon to be 1 and some to be 0). We also enforce that exactly one codon is chosen for each sequence *n* and amino acid position *p* as follows

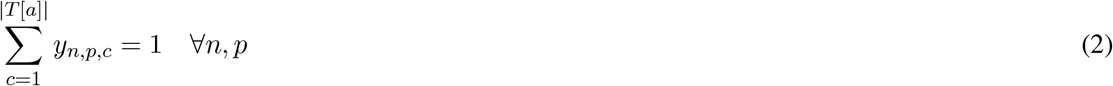

### Auxiliary Variables for Identical Bases

To model common subsequences of increasing length, we first define auxiliary variables representing single identical bases in each base *b* of codon *c* for each amino acid position *p* across every gene pair. For every unordered pair of genes (*s, t*), for each position *p*, codon index *c*, and base index *b*, we define an auxiliary binary variable *z*_*s,t,p,b,c*_ which is set to one if the nucleotide at (*s, p, b, c*) and the nucleotide at (*t, p, b, c*) are identical. To do so, we establish the following relation between *z*_*s,t,p,b,c*_ and the base variables *x*_*s,p,b,c*_ and *x*_*t,p,b,c*_

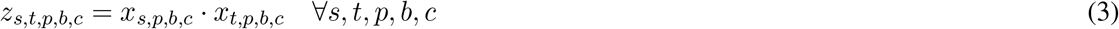

This equation links identical bases in the identical codons for amino acid *p* between the sequence pair *s* and *t*. Now, we want to expand this relationship so that the codon selector can vary for each base column *b*. Formally, for each pair of sequences (*s, t*), amino acid position *p*, and base position *b*, if codons *c* (in gene *s*) and *c*^*′*^ (in gene *t*) have the same base at column *b*, we define a binary variable *z*_*s,t,p,b,c,c*_*′* as follows

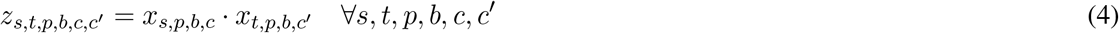

To maintain a compact ILP, we do not introduce the variable *z* when the bases disagree. We note that only one *z* can be set to 1 at a given time because only one codon may be chosen for a given amino acid position. Thus, we can aggregate these auxiliary variables as follows

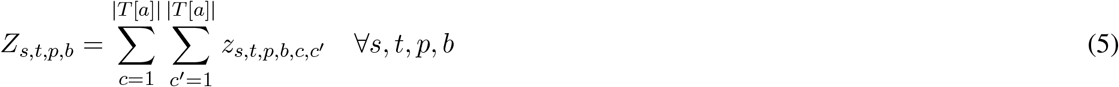

### Modeling Common Subsequences

Observe that *Z*_*s,t,p,b*_ = 1 if gene *s* and gene *t* have the same base in position *b* for amino acid *p*, i.e., *Z*_*s,t,p,b*_ = 1 if *s* and *t* share a common subsequence of length one at position 3*p* + *b*. To simplify the notation, we will drop *s* and *t* (when it is implied) and replace *p* and *b* with position *i* = 3*p* + *b* in the gene sequence. Thus, *Z*_*s,t,p,b*_ = *Z*_*s,t,i*_ = *Z*_*i*_ = 1 when *s*[*i*] = *t*[*i*].

Multiplying the *Z*_*i*_ variables together allows us to represent longer common subsequences. Given two DNA sequences *s* and *t*, observe that *Z*_1_ · *Z*_2_ · *Z*_3_ · · · · · · ·*Z*_*k*_ = 1 if and only if the bases at positions [1, 2, …, *k*] in string *s* match exactly the corresponding bases in string *t*, i.e., there is a shared subsequence of length *k* between *s* and *t* starting at position 1. Similarly, observe that *Z*_2_ · *Z*_3_ · · · · · · ·*Z*_*k*+1_ = 1 if and only if the symbols at positions [2, …, *k* + 1] in *s* match exactly the corresponding symbols in *t*, i.e., there is a shared subsequence of length *k* between *s* and *t* starting at position 2. In general, we have 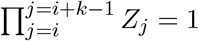 if and only if there is a shared subsequence of length *k* between *s* and *t* starting at position *i*, where 1 ≤ *i* ≤ (3|*P* | − *k* + 1).

### Objective Function

The objective function introduced in section “Problem Definition” requires minimizing pairwise common subsequences in *lexicographic* order, i.e., starting from the longest common subsequence and, in the case of a tie, minimizing progressively shorter common subsequences. Given the maximum length *k* of common subsequences that we want to consider, the solver first determines a solution that minimizes the number of subsequences of length *l* = *k*, recording all variable assignments and objective values. In the next round, the solver minimizes subsequences of length *l* − 1, while enforcing constraints to prevent changes to the previous solution. This process is repeated iteratively until *l* = 1, at which point the final solution is reported. In practice, however, this lexicographic approach is prohibitively slow. We thus adopted a single additive objective function in which the solver minimizes the cumulative contribution of common subsequences across all lengths. The total length of all common subsequences up to length *k* between gene *s* and gene *t* is 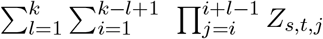. Thus, we minimize over all pairwise sequences (*s, t*) with *s < t* as follows

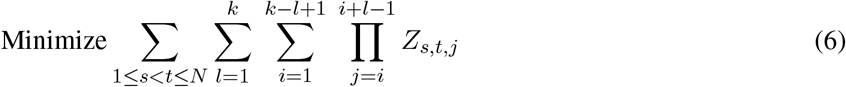

### Solving the Integer Linear Program

Given a protein sequence *P* and the number *N* of desired genes, SIRIUS generates an integer linear program where the binary variables are associated with codon choices and variable-length common subsequences. The constraints of the ILP enforce correct translation and host-specific codon-usage thresholds. SIRIUS solves the ILP with the Google OR-Tools CP-SAT solver [13]. For realistic inputs where |*P*| ≈ 300 and *N* = 10, the resulting ILP has several million binary variables and constraints, thus solving it directly is slow.

To improve the computation time, we use a *warm-start* heuristic [14, 15, 16]. A warm-start heuristic is a hybrid optimization strategy that combines a fast (but suboptimal) greedy solution with a robust exact solver. The goal is to accelerate convergence by supplying the ILP solver with a feasible initial solution, thus reducing the time required to reach convergence. SIRIUS first generates *N* gene sequences with GeneDiversifier [12] using their suboptimal greedy construction, then uses these sequences as starting assignments for all base variables. While these initial assignments provide a good (but suboptimal) feasible solution, the solver remains free to reassign values to base variables to obtain a better solution. The warm-start solution also provides the length of the longest common subsequence observed across any sequence pairs, which determines the value of *k* in the objective function. SIRIUS then optimizes an objective that breaks down these longest subsequences (if possible), followed by the remaining shorter subsequences. This heuristic substantially accelerates convergence while preserving feasibility with respect to codon constraints.

### Filtering Host-Disfavored Codons

An optional feature of SIRIUS allows users to exclude codons that are known to be disfavored in the host organisms, either deterministically or probabilistically. Codon bias is commonly quantified by *relative synonymous codon usage* (RSCU), which captures over- or under-representation of codons across the host’s transcriptome. Users may set a hard RSCU threshold to eliminate disfavored codons before constructing the ILP. However, since rare codons can still be translated at low rates, permitting limited use of such codons can expand the design space and facilitate greater sequence divergence. To this end, users may alternatively set a soft RSCU threshold *τ*_soft_ that allows SIRIUS to choose codons probabilistically. SIRIUS uses the following probability distribution to choose codon *c*

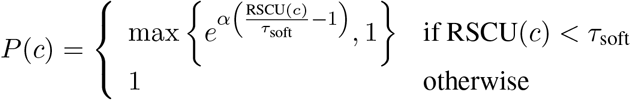

where *α* controls the sharpness of the penalty; smaller RSCU values yield lower inclusion probabilities, while codons above the soft threshold are not penalized.

As an additional safeguard against the overuse of low-RSCU codons, SIRIUS enforces a configurable cap on their frequency. For any codon *c* with RSCU(*c*) *< τ*_soft_, its usage is limited to at most a fraction *ρ* of the positions at which its amino acid occurs, where *ρ* is a user-adjustable value ∈ (0, 1] (e.g., *ρ* = 0.3 for 30%). The constraint is applied independently for each gene sequence and each amino acid. If an amino acid occurs *L* times, a low-RSCU codon for that amino acid may be used no more than ⌊*ρL*⌋ times, regardless of probabilistic sampling. Overall, the mechanisms described in this section aim to accommodate host preference while retaining occasional low-frequency codons to improve combinatorial diversity.

## Results

To reiterate, the inputs to SIRIUS are (a) a protein sequence *P* and (b) the desired number *N* of synonymous genes that translate into *P* . We executed SIRIUS on seven proteins commonly used in biomanufacturing bacterial strains, namely (1) mCitrine fluorescent protein, (2) interferon alpha-2 (*IFNA2*), (3) colony-stimulating factor 3 (*CSF3*), (4) erythropoietin (*EPO*), (5) tissue-type plasminogen activator (*PLAT*), (6) insulin-like growth factor 1 (*IGF1*), and (7) *P. antarctica* lipase B (*PALB/CALB*) [17, 18, 19, 20, 21, 22, 23, 24, 25, 26, 27]. Additional descriptions for these proteins are provided in **Supplementary Table 1**.

We chose *N* = 10 as the number of copies, following previous studies [12, 20]. Although the optimal gene copy number is host- and gene-dependent [28], we selected *N* = 10 to demonstrate SIRIUS’ ability to design synonymous sequences under a realistic multi-copy expression scenario. SIRIUS also allows users to optionally provide a codon usage table tailored to their organism of interest.

The coding sequences generated by SIRIUS for the seven proteins of interest are provided in **Supplementary Data 1**. All experiments were carried out on a high-performance computing (HPC) cluster due to the memory requirements for the ILP solver.

In the following experimental results, we employed GeneDiversifier as both (i) a warm-start generator to accelerate the solving of the ILP and (ii) a method for performance comparison. In the warm-start experiments, the output sequences of GeneDiversifier were provided to SIRIUS to initialize the base variables; in the benchmark comparisons, we evaluated the sequences produced by GeneDiversifier directly, without SIRIUS’ refinement. To establish a baseline, we also included the results of the “Random method” that chooses codons for a residue randomly with uniform probability. We did not use any species-specific codon usage tables in these experiments. A comparison with CodonTransformer was not possible because CodonTransformer is pretrained and fine-tuned to the codon usage biases of specific organisms. CodonTransformer cannot be run without organism codon biases, limiting its applicability and benchmarking outside those contexts.

### SIRIUS Minimizes Homologous Subsequences in Synonymous Gene Design

In the first set of experiments, we used SIRIUS to generate ten synonymous genes for mCitrine. We ran SIRIUS under time limits of {5, 10, 20, 40, 80, 160, 320, 640} minutes, repeating each experiment three times to capture the variance between runs. Figure 2A reports the distribution of shared subsequence counts across output sequence pairs for lengths longer than 10, 12, and 14 bases. Observe that allowing the solver more time improves the quality of the solution by reducing both the number and length of shared subsequences. Specifically, the number of shared subsequences of at least 10 bases decreased by 15.6% when the solver was allowed to run for 80 minutes compared to 5 minutes. Extending the runtime further to 640 minutes yielded diminishing returns, improving the solution by only an additional 1.2% (a total reduction of 16.8% relative to 5 minutes). Based on these results, we selected 80 minutes as the default runtime limit for all subsequent analyses. We recorded the amount of main memory (RAM) required to run SIRIUS as a function of *N* in **Supplementary Figure 1**.

**Figure 2:**
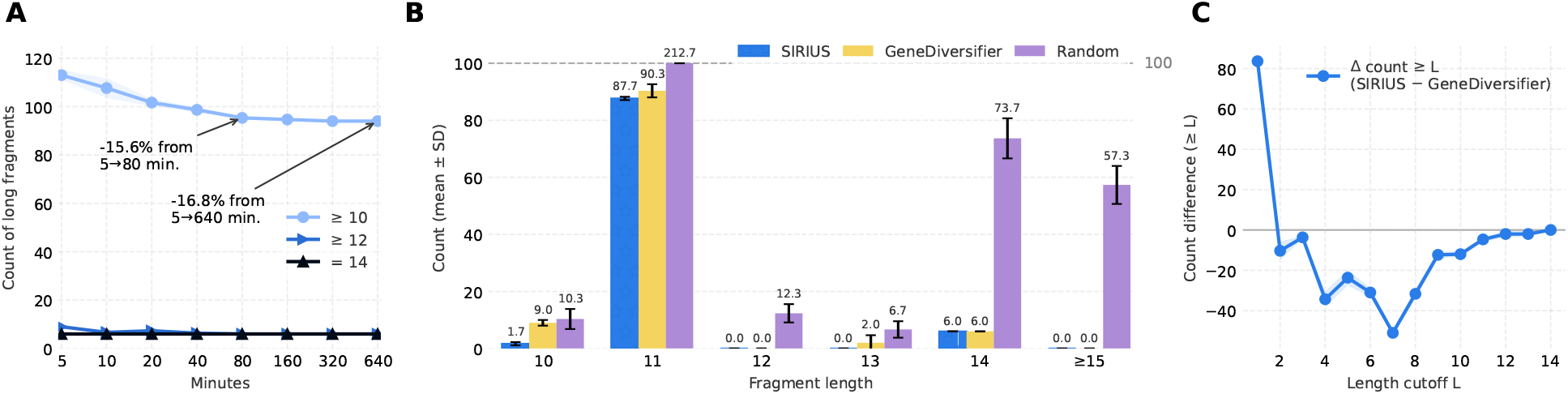
SIRIUS reduces the number and length of common subsequences. . **(A)** Counts of common subsequences longer than 10, 12, and 14 nucleotides as a function of the time allowed for the solver; values are the mean across three replicates; shaded bands show the standard deviation from the mean; **(B)** Mean and standard deviation of counts of common subsequences longer than 10, 12, 14 and ≥ 15 nucleotides for SIRIUS, GeneDiversifier and the method that assigns codons randomly; bars are capped at 100 and labeled with mean values; **(C)** Mean differences between the counts of common subsequences longer than *L* bases between the solution generated by SIRIUS and the solution produced by GeneDiversifier.

Next, we compared the outputs of SIRIUS, GeneDiversifier, and the random method when instructed to generate 10 synonymous genes for mCitrine. We ran each tool three times to capture the variance between executions because two of these methods are non-deterministic. Figure 2B shows the number of pairwise shared subsequences of length 10 − 14 (and longer than 15) nucleotides for the three methods. Better solutions have fewer occurrences of long shared subsequences. However, breaking longer subsequences into smaller subsequences can increase the number of shorter subsequences. Observe that SIRIUS solutions have fewer shared subsequences of length 13 and 11 nucleotides than GeneDiversifier. Notably, for subsequences of length 10 nucleotides, SIRIUS produces an average of 1.7 shared subsequences across three runs, compared to 10 for GeneDiversifier. Although SIRIUS exhibits higher counts at shorter subsequence lengths (9, 6, 3, and 1 nucleotides), these lengths are unlikely to drive homologous recombination [29, 30]. Observe that neither SIRIUS nor GeneDiversifier can eliminate the 6 common subsequences of length 14. This limitation does not reflect a shortcoming of the algorithms but rather the constraints of the genetic code: beyond a certain subsequence length, the number of synonymous codon combinations is insufficient to avoid repeats entirely, making these long subsequences unavoidable. Neither algorithm produces common subsequences longer than 14 nucleotides, while the random approach can output shared subsequences of up to 57 nucleotides.

We provide the same comparative analysis for the remaining six proteins in **Supplementary Figures 2–7**. For example, for genes encoding the protein *CALB*, SIRIUS generates sequences with fewer common subsequences of length 13 compared to GeneDiversifier (0.0 vs. 0.7), 12 (0.0 vs. 1.7), 11 (49.3 vs. 50.0), and notably 10 (1.3 vs. 8.0), averaged across three runs per tool (**Supplementary Figure 2**).

In Figure 2C, we compared SIRIUS and GeneDiversifier by plotting

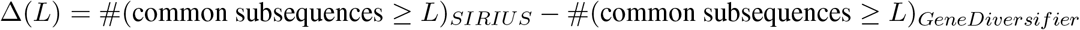

for various choices of the thresholds *L*, averaging over three independent runs to capture stochastic variations. Observe that after *L* ≥ 2 this difference is always negative, reaching a minimum at *L* = 7 (51 less common subsequences produced by SIRIUS compared to GeneDiversifier). This indicates that SIRIUS can produce solutions with much less common subsequences than GeneDiversifier.

## Conclusions

In this work, we introduced SIRIUS, a novel design tool capable of producing maximally divergent DNA sequences for a given target protein. SIRIUS leverages integer linear programming to generate synonymous sequences with the fewest possible pairwise common subsequences. Previous approaches have relied on heuristics, machine learning, or other suboptimal algorithms. Our experiments show that while these methods can make local improvements, they struggle to capture the full combinatorial context of the problem and therefore often yield suboptimal solutions.

As a proof of concept, we applied SIRIUS to design divergent sequences for seven therapeutically and industrially relevant proteins. Because solving an ILP with millions of variables and constraints is computationally intractable, we leveraged GeneDiversifier as a warm-start to initialize the solver. This strategy drastically reduced the runtime and enabled SIRIUS to consistently improve upon GeneDiversifier’s solutions.

While SIRIUS overcomes key algorithmic limitations of previous heuristic-based methods, a major challenge remains its substantial memory and runtime requirements, particularly for longer proteins and larger values of *N* . We anticipate that introducing conditional constraints tailored to the structure of the problem will be critical for reducing the ILP complexity and improving scalability. Nonetheless, SIRIUS provides a novel and effective framework for designing maximally divergent coding sequences, enabling more stable multi-copy expression in biomanufacturing strains.

## Data Availability

SIRIUS is freely available on GitHub at https://github.com/ucrbioinfo/sirius and on Zenodo at 10.5281/zenodo.18343092. The documentation for SIRIUS is provided on its GitHub Wiki page. All data used in this study can be obtained from the GitHub and Zenodo repositories and the **Supplementary Data**.

## Supplementary Data

Supplementary Data are available.

## Author Contributions

**Amirsadra Mohseni**: Conceptualization, Software, Methodology, Formal Analysis, Data Curation, Visualization, Writing – Original Draft. **Stefano Lonardi**: Conceptualization, Supervision, Project Administration, Funding Acquisition, Writing – Original Draft. **Ian Wheeldon**: Conceptualization, Funding Acquisition, Writing – Original Draft.

## Acknowledgements

We thank the developers of Google OR-Tools, GeneDiversifier, and the Non-Repetitive Parts Calculator for making their software openly available.

## Footnotes

* In German, *Systematische Identifikation Redundanter Identisch Übersetzter Sequenzen*.

