## Supplementary Material for "Maximally Divergent Synonymous Gene Design with SIRIUS"

### Maximally Divergent Synonymous Gene Design with SIRIUS (Supplementary Material)

#### Supplementary Notes

In this work, we apply the following algorithms to quantify the quality of a solution output by GeneDiversifier, SIRIUS, and other tools considered. An effective tool is one that lexicographically minimizes the lengths of identical contiguous runs, and when tied in lengths, reduces the number of their occurrences. Below is a pseudocode reflecting the internal functions employed. Note that Algorithm 2 on page 2 calls Algorithm 1.

---

**Algorithm 1** Detection of contiguous exact-match segments in one aligned pair. Given two equal-length sequences, scan left to right and record each gap-free run of identical characters as a triple (start,end,length). Only positionwise matches are considered; runs end at the first mismatch or the sequence end.

---

**Input:** Two equal-length sequences X and Y of length $L$
**Output:** List of runs as triples (start_index, end_index, length)

```
procedure FindHomologousRuns(X, Y)
    runs ← empty list
    start ← NONE
    for i ← 0 to L − 1 do
        if X[i] = Y[i] then
            if start = NONE then
                start ← i
            end if
        else
            if start ≠ NONE then
                Append (start, i − 1, i − start) to runs
                start ← NONE
            end if
        end if
    end for
    if start ≠ NONE then
        Append (start, L − 1, L − start) to runs
    end if
    return runs
end procedure
```

**Algorithm 2** Distribution of run lengths across all pairs. For a collection of equal-length aligned sequences, apply Algorithm 1 to every unordered pair and tally segment lengths to obtain the run-length frequency distribution, optionally reported from longest to shortest.

**Input:** Collection S of $N$ equal-length sequences (aligned)
**Output:** Frequency map F from run length $\rightarrow$ count

**procedure** FINDALLHOMOLOGOUSRUNSANDCOUNTLENGTHS(list of sequences S)
  F $\leftarrow$ empty map with default value 0
  **for** each unordered pair $(a, b)$ with $0 \leq a < b < N$ **do**
    R $\leftarrow$ runs from Algorithm 1 applied to $S[a]$ and $S[b]$
    **for** each triple (_, _, len) in R **do**
      F[len] $\leftarrow$ F[len] $+1$
    **end for**
  **end for**
  **return** F (reported sorted by decreasing length)
**end procedure**

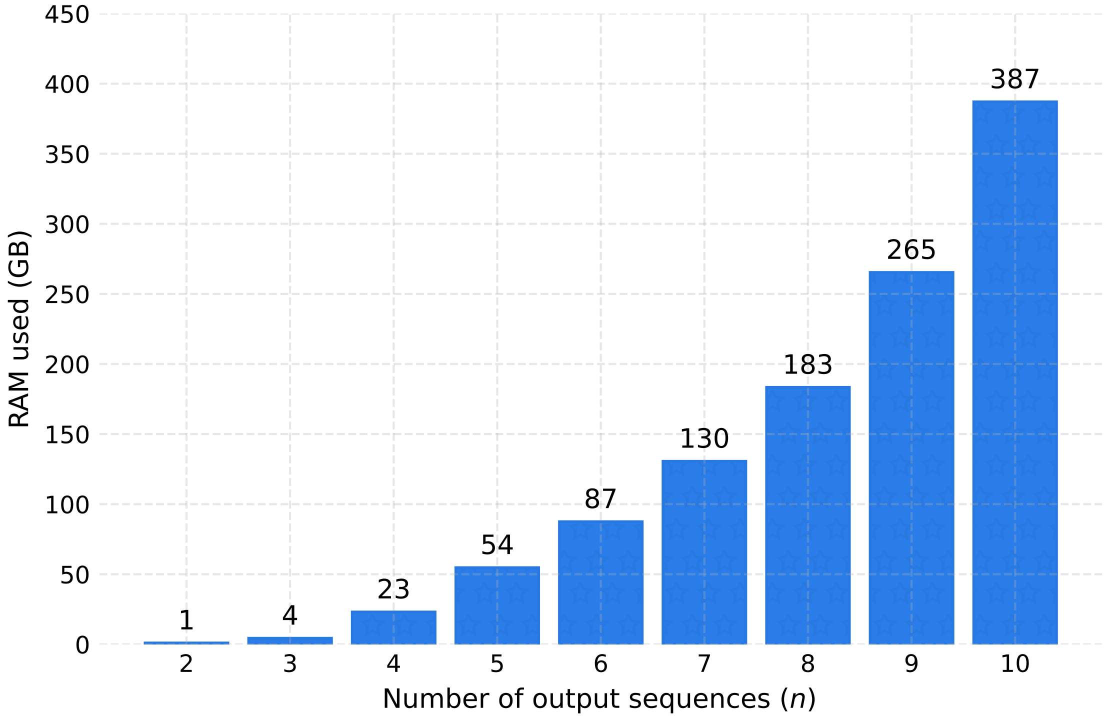


**Supplementary Figure 1**. Main (RAM) memory usage of SIRIUS for $P$ = mCitrine and $n \in [1, 10]$ rounded up to the nearest GB.

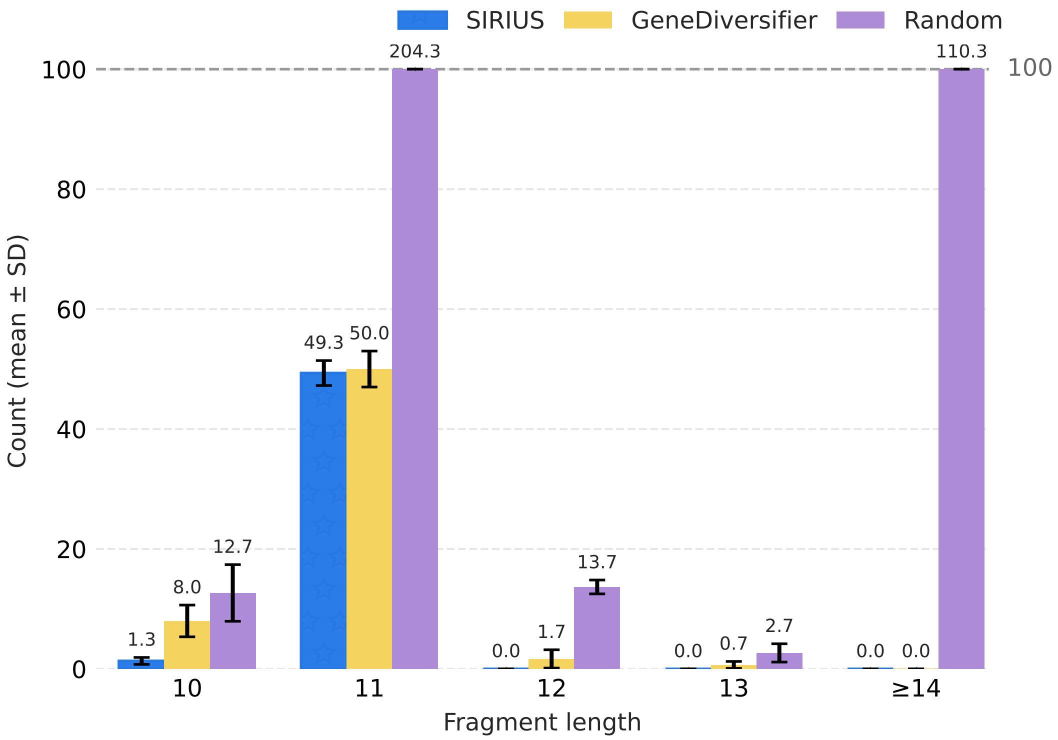


**Supplementary Figure 2**. Using the *CALB* protein and $N = 10$ as input, this figure shows the mean and standard deviation of counts of common subsequences longer than 10 nucleotides for SIRIUS, GeneDiversifier and the random method; bars are capped at 100 and labeled with mean values across three runs for each tool.

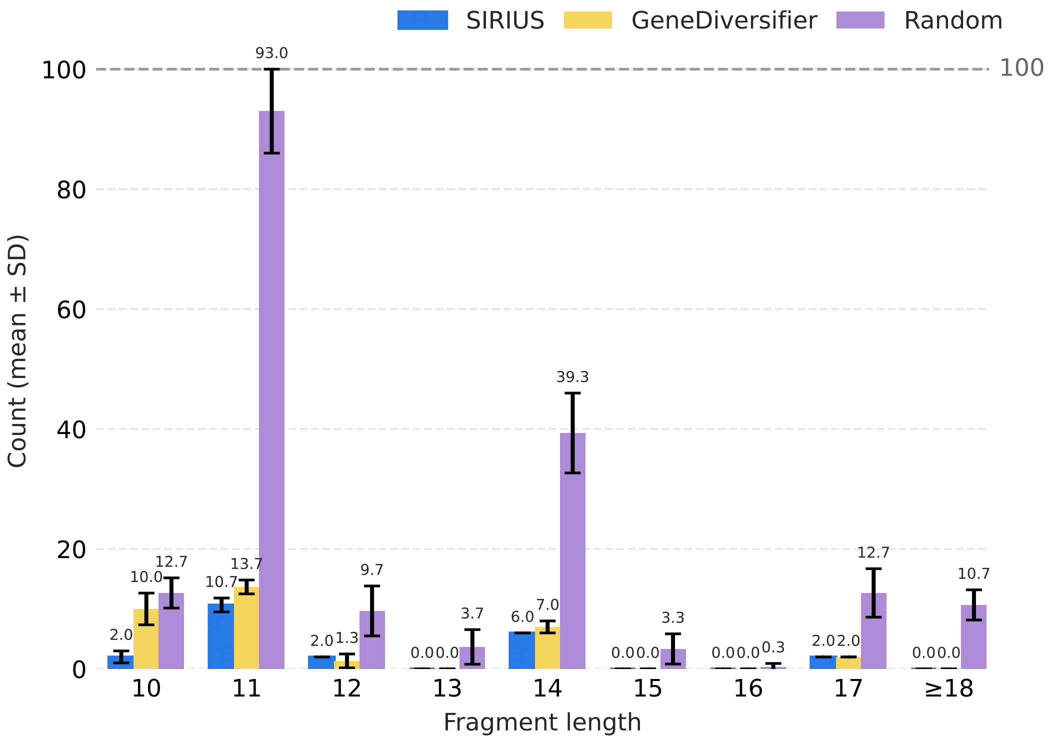


**Supplementary Figure 3**. Using the *CSF3* protein and $N = 10$ as input, this figure shows the mean and standard deviation of counts of common subsequences longer than 10 nucleotides for SIRIUS, GeneDiversifier and the random method; bars are capped at 100 and labeled with mean values across three runs for each tool.

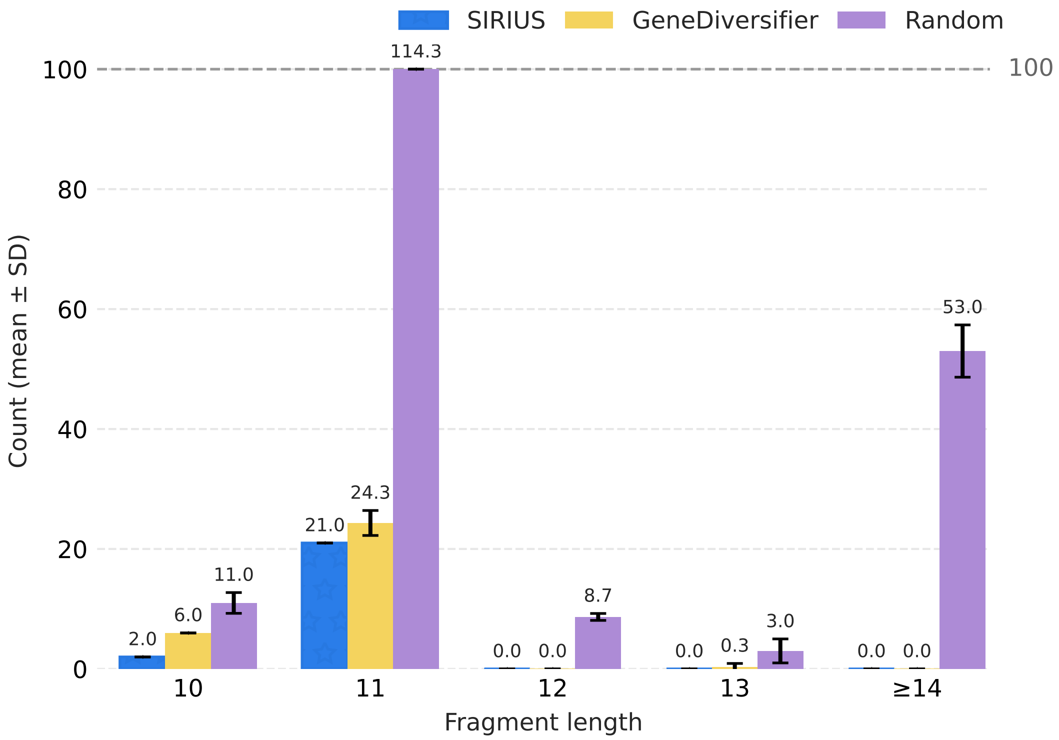


**Supplementary Figure 4**. Using the *EPO* protein and $N = 10$ as input, this figure shows the mean and standard deviation of counts of common subsequences longer than 10 nucleotides for SIRIUS, GeneDiversifier and the random method; bars are capped at 100 and labeled with mean values across three runs for each tool.

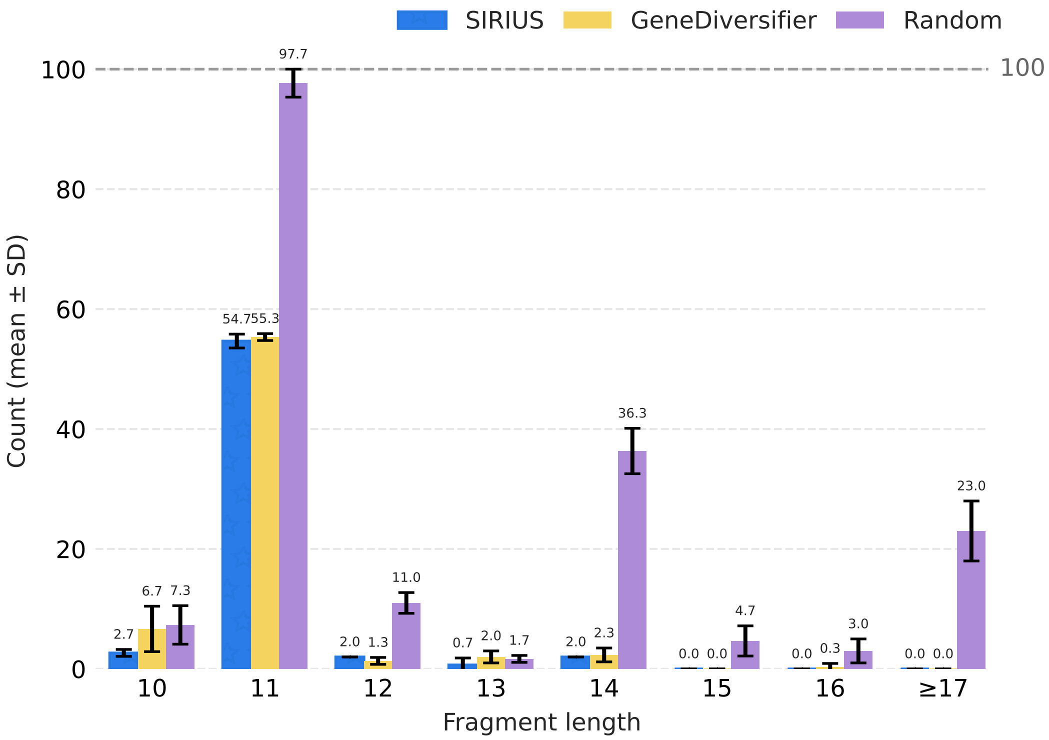


**Supplementary Figure 5**. Using the *IGF1* protein and $N = 10$ as input, this figure shows the mean and standard deviation of counts of common subsequences longer than 10 nucleotides for SIRIUS, GeneDiversifier and the random method; bars are capped at 100 and labeled with mean values across three runs for each tool.

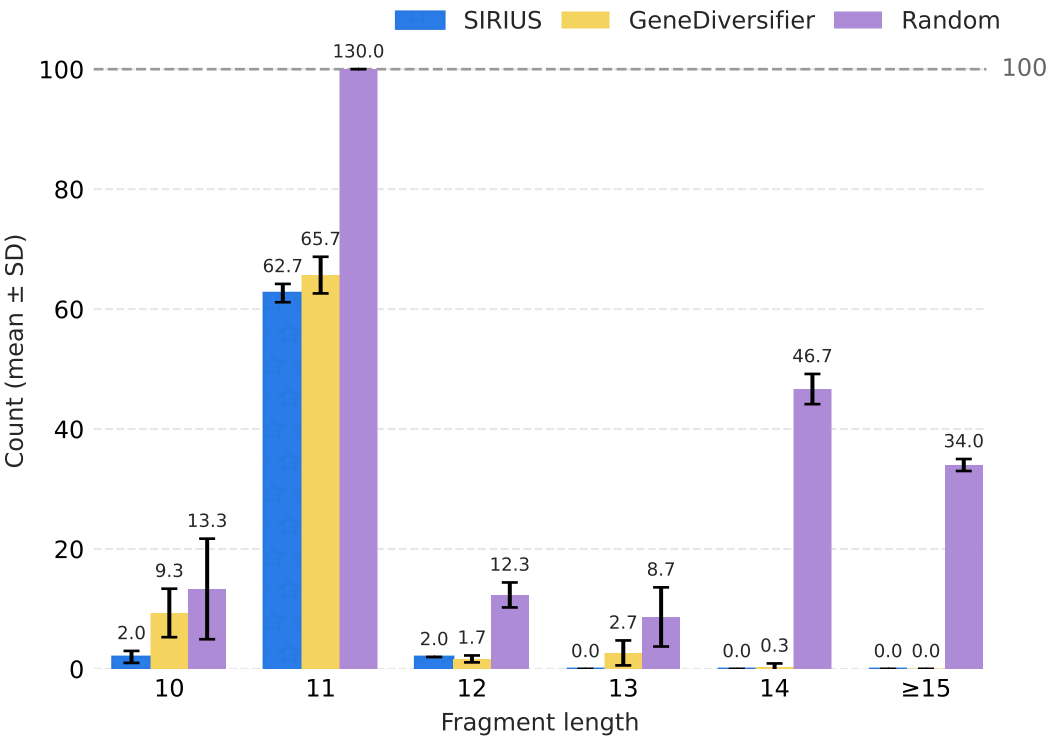


**Supplementary Figure 6**. Using the *INFA2* protein and $N = 10$ as input, this figure shows the mean and standard deviation of counts of common subsequences longer than 10 nucleotides for SIRIUS, GeneDiversifier and the random method; bars are capped at 100 and labeled with mean values across three runs for each tool.

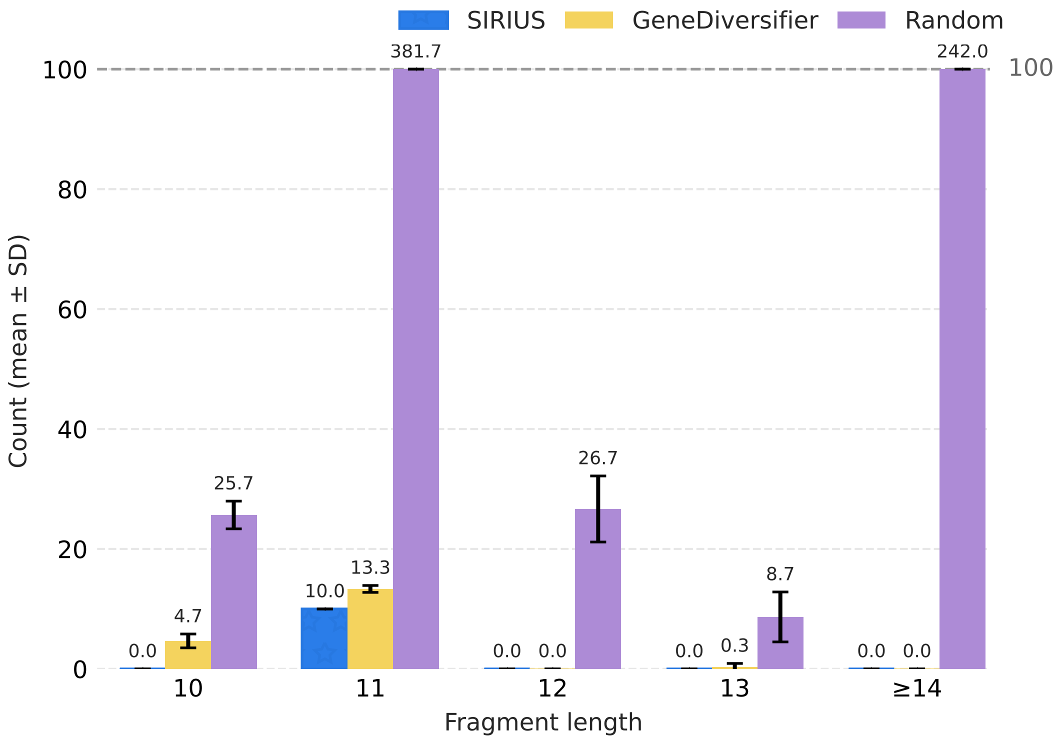


**Supplementary Figure 7**. Using the *PLAT* protein and $N = 6$ as input, this figure shows the mean and standard deviation of counts of common subsequences longer than 10 nucleotides for SIRIUS, GeneDiversifier and the random method; bars are capped at 100 and labeled with mean values across three runs for each tool. Due to the intensive RAM demands of this instance, a smaller value for $N$ was chosen.

**Supplementary Table 1.** Brief functional description of the seven pharmaceutical- and industrially relevant genes.

| Gene | Description | Common Expression Host |
|---|---|---|
| *IFNA2* | This gene is a member of the alpha interferon gene cluster on chromosome 9. The encoded cytokine is a member of the type I interferon family that is produced in response to viral infection as a key part of the innate immune response with potent antiviral, antiproliferative and immunomodulatory properties [1]. | *E. coli* [2], *P. pastoris* [3] |
| *CSF3* | Encodes a member of the IL-6 superfamily of cytokines. The encoded cytokine controls the production, differentiation, and function of granulocytes. A modified form of this protein is commonly administered to manage chemotherapy-induced neutropenia [4]. | *E. coli* [5], CHO cells [6, 7] |
| *EPO* | Encodes a secreted, glycosylated cytokine mainly produced in the kidney and released into the bloodstream. It binds to the erythropoietin receptor to stimulate red blood cell formation in the bone marrow. Recombinant versions also show neuroprotective and anti-apoptotic effects in various tissues and are used to treat anemia and enhance cancer therapy outcomes [8]. | HEK293 and CHO cells [9] |
| *PLAT/ TPA/ t-PA* | Encodes a secreted serine protease that converts plasminogen into plasmin, an enzyme responsible for breaking down blood clots. The protein is synthesized as a preproprotein that is cleaved into heavy and light chains, which form a disulfide-linked heterodimer. It functions in fibrinolysis, cell migration, and tissue remodeling, and recombinant forms are FDA-approved for treating acute ischemic stroke [10, 11]. | *E. coli* [12], CHO cells [13], *P. pastoris* [14] |
| *IGF1* | The protein encoded by this gene is similar to insulin in function and structure and is a member of a family of proteins involved in mediating growth and development. This gene can be expressed in biomanufacturing hosts such as *P. pastoris* IGF1 and has potential in the treatment of a variety of neuronal disorders and injuries [15]. | E. coli [16], *P. pastoris* [17] |
| *PALB/ CALB* | Lipase B from *Pseudozyma (Candida) antarctica* is a versatile and stable biocatalyst used in laboratory and industrial organic synthesis. It exhibits high enantioselectivity, thermal stability, broad substrate specificity, and solvent tolerance, enabling applications such as polymerization, kinetic resolution of alcohols and amines, β-lactam ring opening, and desymmetrization of complex drug intermediates [18]. | *E. coli* [19], *P. pastoris* [20] |
| mCitrine | mCitrine is a basic (constitutively fluorescent) yellow fluorescent protein. mCitrine is not a naturally occurring protein; it is a genetically engineered monomeric YFP that was derived from the original Green Fluorescent Protein (GFP) found in the *A. victoria* jellyfish [21]. | Variable |
